# Hemoglobin gene repertoire in teleost and cichlid fishes shaped by gene duplications and genome rearrangements

**DOI:** 10.1101/2024.03.26.586788

**Authors:** Dmytro Omelchenko, Arnold Roger Bitja-Nyom, Michael Matschiner, Milan Malinsky, Adrian Indermaur, Walter Salzburger, Oldřich Bartoš, Zuzana Musilova

## Abstract

Hemoglobin is a crucial element of the oxygen transport system in vertebrates. It exhibits remarkable gene diversity across teleost fishes, reflecting their evolutionary adaptations for thriving in various aquatic environments. In this study, we present the dynamic evolution of hemoglobin subunit genes based on a comparison of high quality long-read genome assemblies of 24 vertebrate species, including 16 teleosts (of which six are cichlids). Our findings indicate that teleost genomes contain between five (fugu) and 43 (salmon) hemoglobin genes, representing the largest hemoglobin gene repertoire among vertebrates. We find evidence that the ancestor of teleosts had at least four Hbα and three or four Hbβ subunit genes, and that the current gene diversity emerged during subsequent teleost radiation, driven primarily by (tandem) gene duplications, genome compaction, and rearrangement dynamics. We provide insights into the genomic organization of hemoglobin clusters, revealing the parallel origin of multiple clusters in tetrapods and in teleosts. Importantly, we show that the presence of paralogous rhbdf1 genes flanking both teleost hemoglobin clusters (LA and MN) supports the hypothesis for the origin of the LA cluster by rearrangement within teleosts, rather than by the teleost specific whole-genome duplication. We specifically focus on cichlid fishes, where adaptation to low oxygen environments has been shown to play roles in species diversification. Our analysis of six cichlid genomes, including the *Pungu maclareni* from crater lake Barombi Mbo, for which we sequenced the representative genome, reveals 18 to 31 copies of the Hb genes, and elevated rates of non-synonymous substitutions compared to other teleosts. Overall, this work facilitates a deeper understanding of how hemoglobin genes contribute to the adaptive and diversification potential of teleosts.

## Introduction

Hemoglobin is an oxygen-binding metalloprotein responsible for oxygen transport in vertebrates (Storz, 2018). In jawed vertebrates (gnathostomes), the hemoglobin molecule has a quaternary structure and consists of four globin polypeptide chains (two Hbα and two Hbβ subunits) attached to the prosthetic heme group (Wells, 1999). This tetrameric structure is considered to be of great functional importance since it provides a mechanism for cooperative oxygen-binding and allosteric regulatory control (Storz, Opazo & Hoffman, 2013). Vertebrates have multiple hemoglobin genes and are known for a developmental switch between juvenile and adult hemoglobins, resulting in most vertebrates expressing different genes during ontogenesis (Chan et al., 1997). This differential expression of distinct hemoglobin isoforms helps to deal with the changing oxygen transport challenges encountered during vertebrate development (Opazo et al., 2013).

Vertebrates have diversified following two rounds of whole-genome duplications in their ancestor (Ohno, 1970), and teleost fishes have undergone an additional teleost-specific genome duplication (TSGD) (Kuraku & Meyer, 2009 and ref therein). The evolutionary success of both vertebrates and teleosts has often been associated with these whole-genome duplications thanks to the sufficient genetic “substrate” for selection to act upon (Glasauer & Neuhauss, 2014). For example, the Hox gene family (Holland et al., 1994), MHC genes (Abi-Rached et al., 2002) or parvalbumin genes (Mukherjee et al., 2021) still carry the traces of the whole-genome duplications. In general, many gene families have been shaped by the interplay between whole-genome duplications, ancestral or lineage-specific (tandem) gene duplications, and prevalent gene losses (Hughes & Freidman, 2003; Rennison et al., 2012; Steinke et al., 2006, Parey et al., 2022), and that includes also teleost hemoglobins (Quinn et al., 2010; Opazo et al., 2013; Storz et al., 2016).

Hemoglobin genes (Hb) originated together with other globin genes (myoglobin, cytoglobin) after two rounds of whole-genome duplications in the vertebrate ancestor (Hoffmann, Opazo & Storz, 2011). Subsequent subfunctionalization of gas transport and oxygen storage between hemoglobin and myoglobin is sometimes seen as a key innovation of early vertebrates (Storz, Opazo & Hoffman, 2013). The Hb subunit genes - proto-Hbα and Hbβ have afterwards resulted from an ancient tandem duplication in early jawed vertebrates (Opazo et al., 2012). Hemoglobin subunit genes are found in multi-copy tandem repeats and the ancestral genomic arrangement was probably a single cluster with Hbα and Hbβ subunit genes mixed, as still seen in cartilaginous fish (Marino et al., 2007). Teleosts have hemoglobins organized in two genomic clusters (labelled as MN and LA), which both carry mixed Hbα and Hbβ subunits, while in tetrapods the Hb genes have been rearranged into a separate Hbα and Hbβ clusters (Hardison, 2008; Patel, et al., 2008). Homologs of the upstream flanking genes of the teleost MN cluster are also found upstream of the Hbα cluster in tetrapods, while the LA cluster seems to be specific for teleosts only (Opazo et al., 2012; Hardison, 2012) and it has been suggested that the LA cluster originated from the teleost specific whole-genome duplication (TSGD) (Opazo et al., 2013). Teleost fishes have a greater variation and often a high number of hemoglobin genes compared to air-breathing vertebrates, which can be explained by the fact that variation in oxygen availability is generally much greater in the aquatic environment (Opazo et al., 2013).

In addition to the use of multiple Hb isoforms, hemoglobins diversify and adapt to variation in oxygen availability by sequence evolution. Several studies have shown signatures of positive selection on the amino acid sites in particular Hb gene copies. Specifically, an excess of non-synonymous substitutions in nucleotide sequence of hemoglobins is found in species from extreme environments, such as in lagomorphs and rodents from Qinghai-Tibet plateau (Chen et al., 2016), or two cetacean species (dolphin and killer whale) where it was hypothesized that it could lead to an adaptation to their prolonged dives (Nery et al., 2013). In teleosts, it has been shown that high-altitude schizothoracine fishes from Qinghai-Tibet plateau have functionally divergent hemoglobin genes with positively selected sites (Lei et al., 2021).

Some teleost families diversify more and occupy much broader range of niches than others. The cichlid fish represent one of the most species-rich families of vertebrates, known for their large and rapid adaptive radiations, especially in the tropical lakes of Africa (Seehausen, 2000; Kautt et al., 2016; Svardal et al., 2021) and serve as a model group for research into adaption and diversification (e.g., Ronco et al., 2021; Musilova et al., 2019). Based on five initial short-read genome assemblies, it has been shown that cichlids have elevated number of gene duplicates (Brawand et al., 2014), but the particular case of hemoglobin genes has not been investigated. Warm tropical lakes, including the lakes hosting cichlid fish radiations, tend to be permanently stratified with steep oxygen gradients. Nevertheless, deep waters with low oxygen concentrations have repeatedly been colonized by cichlids, and depth adaptation represents one of the most important axes of differentiation in multiple lakes, e.g., Lake Malawi (Malinsky et al., 2018), Lake Masoko (Malinsky et al., 2015), Lake Barombi Mbo (Green et al., 1973; Musilova et al., 2019), or Lake Tanganyika (Ronco et al., 2021). It has been shown that deep-water adapted cichlid species from Barombi Mbo have highly elevated blood hemoglobin concentrations (Green et al., 1973) and that six hemoglobin genes contain signs of parallel adaptation in the Lake Malawi deep-water cichlids groups *Diplotaxodon* and ‘deep-benthic’ (Hahn et al., 2017; Malinsky et al., 2018). However, a more comprehensive analysis of selection on hemoglobin genes across multiple cichlid adaptive radiations is lacking.

Hemoglobin gene clusters are highly repetitive and correct understanding of their evolutionary dynamics requires accurate genome assemblies. Here we reconstruct the evolution of the hemoglobin subunit genes based on high quality long-read assemblies of 24 vertebrate species, including 16 teleosts, six of which belong to the cichlid family. Compared with previous analyses, that were based primarily on short read genome assemblies (e.g., Opazo et al, 2013), we find much expanded Hb repertoires, driven primarily by (tandem) gene duplications. Our reconstruction of Hb cluster evolution reveals dynamics of duplication, rearrangements, and genome compaction, and provides evidence that the teleost LA and MN clusters originated after TSGD. We used the comprehensive collection of Hb genes from all 24 genomes to conduct a comprehensive search for signatures of positive selection on the sequence level, revealing a 27 Hbα and 17 Hbβ under positive selection within the cichlid family. Overall, these findings fill in numerous missing puzzles in the picture of dynamic evolution of Hb genes in teleosts, and provide a framework within which we can investigate specific cases of hemoglobin and oxygen transport adaptations, such as in the lake Barombi Mbo.

## Material and methods

### Sampling and DNA extraction

Live specimens were collected in the Barombi Mbo crater lake and transported to the aquarium facility at Charles University in Prague (collecting permits: 0000032,48,49/MINRESI/B00/C00/C10/C12). One male individual of *Pungu maclareni* was euthanized and liver and spleen tissues were dissected for the DNA extraction. In order to obtain High Molecular Weight (HMW) DNA, we homogenized the tissues under liquid nitrogen followed by a standard phenol-chloroform DNA extraction (e.g. McKiernan & Danielson, 2017). DNA was then quantified using a Qubit fluorometer (Invitrogene) and checked for degradation by standard electrophoresis.

Libraries for Oxford Nanopore Technologies (ONT) sequencing were prepared from a HMW DNA using a Ligation Sequencing Kit (SQK-LSK109). Libraries were sequenced on the ONT GridIONx5 platform using the R9.4.1 chemistry (Flow-Cell). Sequencing data were base-called, i.e. transmission from physical changes in the electric current signal measured by the ONT sequencing device to biologically relevant bases, using Guppy v3.2.10 in “high-quality” settings/mode (e.g. Wick et al., 2019).

### Nanopore sequencing and assembly

Basic characteristics of the ONT libraries were estimated in the R 3.6.0 using the NanoR library (R Core Team 2019; Bolognini et al., 2019). In summary, we acquired more than five million reads with an average length of over 7000bp, which in total slightly exceeded 40 Gbp (only reads > 1000 bp were considered; see Supplementary Table 2). The genome was assembled using Flye 2.7.1 long-read assembler (Kolomogorov et al., 2019). The primary assembly was then polished with the ONT long-reads alone using the Medaka 1.0.1 software tool (Lee et al. 2021). The assembly was further iteratively (four times) polished by Whole Genome Sequencing (WGS) Illumina reads using the Pilon software tool (Walker et al. 2014). The quality (completeness) of the assembly was assessed using BUSCO 4.0.6 (actinopterygii_odb10; see Supplementary Table 3) (Simão et al. 2015). The resulting genome sequences were deposited into NCBI GenBank repositories (accession number: *tba*).

### Gene mining and gene annotation

We used our genomic data (*Pungu maclareni*) complemented by a set of selected species with high-quality assemblies from GenBank (see Table 1 for accession numbers). Our focus was on a comparative analysis of the hemoglobin subunit genes repertoire and its genomic organization. In total, we analyzed 24 genomes, 16 of teleosts and six of other vertebrates (see Table 1). We cover all four main teleost lineages (i.e., Elopomorpha, Osteoglossomorpha, Otomorpha and Euteleostei) and specifically focus on cichlids. We used following bioinformatic tools to mine for hemoglobin genes. Geneious software ver. 9.1 with High sensitivity setting (http://www.geneious.com, Kearse et al., 2012) was used to capture scaffolds/chromosomes carrying Hb genes by mapping the assembled genome against the zebrafish reference composed of one Hbα (NM_131257.3) and one Hbβ gene exons (BC139602.1). If an annotation was available, we extracted all hemoglobin genes from the respective genome. We subsequently manually mapped the single exons of all extracted genes against flanking regions of the identified genes and also to the intergenic regions to reveal the entire Hb gene cluster and confirmed the annotation using the Geneious software. In some cases, this approach served to improve the annotation. Furthermore, we performed AUGUSTUS *ab initio* gene prediction (Stanke & Morgenstern, 2005) to reveal hemoglobin genes in the unannotated eel genome (*Anguilla anguilla*) with a Nanopore assembly (accession number PRJEB20018 in the European Nucleotide Archive). We inspected the *Pungu maclareni* genome and we have manually annotated the found Hbα and Hbβ genes.

### Synteny analysis

We manually inspected genomic regions that contain hemoglobin subunits genes (both MN and LA clusters) in five cichlid species with an assembled genome in a public access: (Nile tilapia (*Oreochromis niloticus*), Eastern happy (*Astatotilapia calliptera*), zebra mbuna (*Maylandia zebra*), Flier cichlid (*Archocentrus centrarchus),* and Midas cichlid (*Amphilophus citrinellus*). Furthermore, we have similarly identified hemoglobin clusters in *Pungu maclareni* based on our manual gene annotation. Genomic sequences of hemoglobin clusters were extracted from the linkage groups including the whole sequence between flanking genes and reoriented for further comparison of synteny with the nprl3 gene upstream to the hemoglobin genes in the MN cluster, and aqp8 upstream to the hemoglobin subunits in the LA cluster. We subsequently used Gepard 1.40 (Krumisek et al., 2007) to create comparative dotplots of the hemoglobin clusters and identified traces of gene duplications or inversions.

### Phylogenetic analysis of the Hb genes

To complement the teleost data set, we included also the hemoglobin clusters of six non-teleost fishes and tetrapods (Table 1, Figure 1). Coding sequences of the hemoglobin genes were extracted and used for the construction of a combined gene tree for both Hbα and Hbβ subunits. Cytoglobins from Australian ghost shark, spotted gar and Nile tilapia were used as an outgroup. The nucleotide sequences of the extracted Hb genes were aligned with MAFFT ver. 7.222 (Katoh & Standley, 2013). The best fitting substitution model (GTR+I+G) was selected by jModeltest2 ver. 2.1.6 (Darriba et al., 2012). We performed both Bayesian inference and Maximum Likelihood phylogenetic analysis. Phylogenetic analyses were performed on the CIPRES Science Gateway portal (Miller et al, 2010) employing MrBayes ver. 3.2.6 (Ronquist et al., 2012) for 20 million generations and 12.5% burnin. We used IQ-TREE (Nguyen et al., 2014) to infer phylogenetic trees by maximum likelihood employing SH-aLRT test.with substitution model selection by ModelFinder (Kalyaanamoorthy et al., 2017) and the ultrafast bootstrap approximation (Hoang et al., 2017) with 1000 replicates.

**Figure 1.**
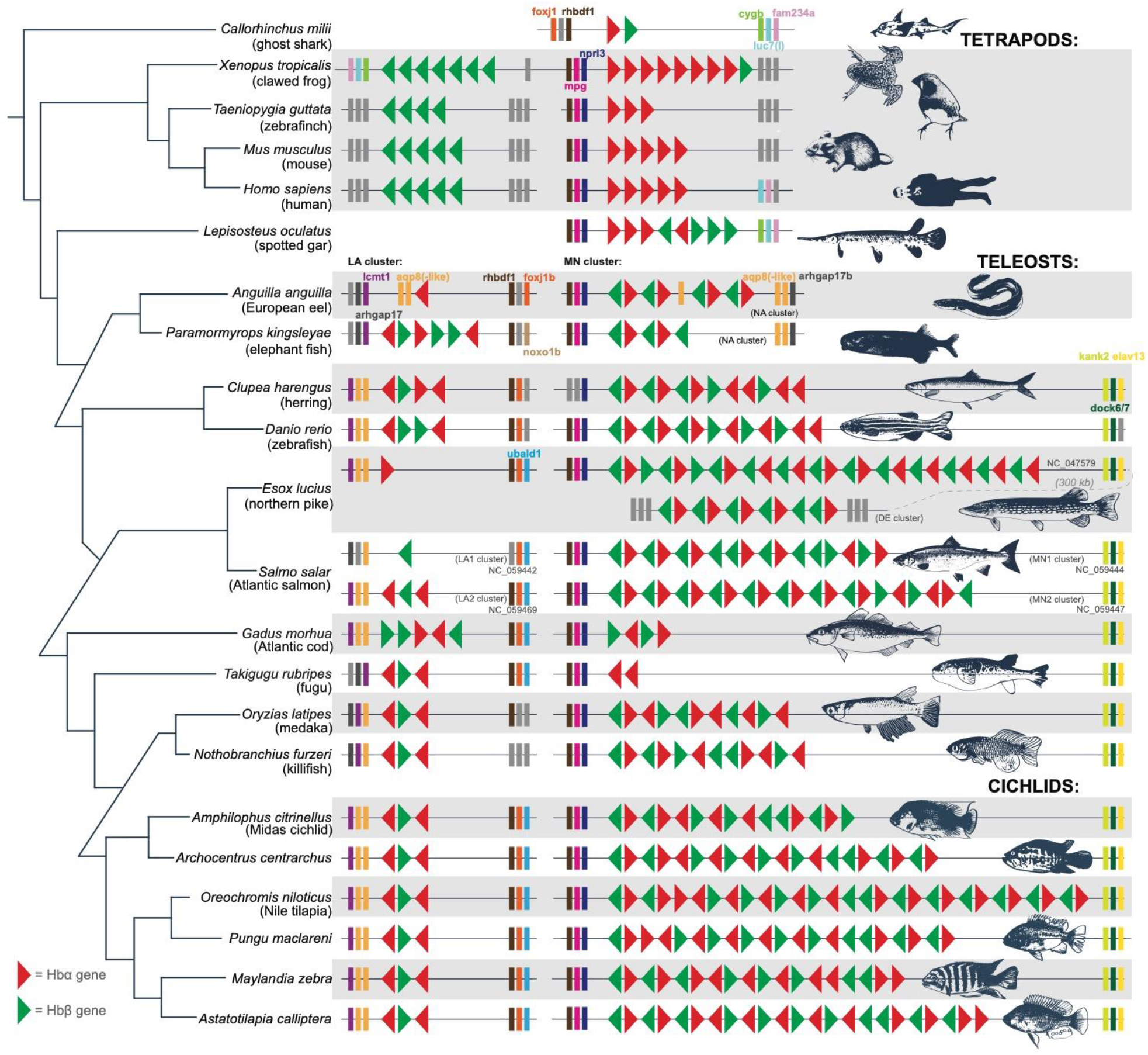
Conserved synteny and genomic organization of the hemoglobin clusters in major linages of gnathostomes, teleosts and cichlids. Hemoglobin genes shown as triangles with the forward or reverse orientation, Hbα subunit genes shown in red, Hbβ subunit genes shown in green. Flanking genes shown as rectangles either having a corresponding color code (for the genes frequent in different evolutionary groups) or in gray if unique. We present the clusters in the same orientation for comparative purposes. The MN cluster refers to a larger and more variable cluster with the mpg and nprl3 upstream flanking genes, whereas the LA cluster is more conserved and named after lcmt1 and aqp-8 flanking genes. Note that in Elopomorpha (eel) and Osteoglossomorpha (*Paramormyrops*) the MN cluster has sometimes been referred to as NA cluster due to the presence of the aquaporin (aqp8) genes. The tree topology is based on relationships after Betancur et al. (2017)

### Positive selection test

The codon alignments were used for the selection test employing CODEML package as a part of the PAML software (Yang, 2007). Hb genes that contained in-frame stop codons were removed from the analysis. We performed a codon alignment in the PAL2NAL software (Suyama et al., 2006). We ran analyses separately for the Hbα and Hbβ subunits to reduce computational time. We used a branch-site model with heterogeneous omega ratios among sites allowed and estimated the ratio between the non-synonymous and synonymous substitutions for background branches. We have subsequently produced final trees with branches colored accordingly to the dN/dS ratio using *treeio* R package (Wang et al., 2020)

## Results

### Hemoglobin gene repertoire and cluster synteny

We analyzed hemoglobin genes in the high-quality genomes of 24 vertebrate species, one cartilaginous fish, four tetrapods, one non-teleost and 16 teleosts, of which six were cichlids (Fig 1). Our cichlid collection included both African and Neotropical cichlids, and to complement the existent cichlid data set we sequenced the representative genome of *Pungu maclareni*, an endemic cichlid species from crater lake Barombi Mbo (Cameroon, Africa) by applying the Oxford Nanopore technology. The assembled genome has a N50 of 3’073’378 with both hemoglobin clusters recovered intact on separate scaffolds.

Our results show that teleost fishes generally have a larger hemoglobin gene repertoire compared to other vertebrates. Teleosts possess between 5 and 43 hemoglobin genes in their genome (Fig. 1). The Japanese pufferfish (*Takifugu rubripes*) has the lowest number of hemoglobin genes (four Hbα subunit genes and one Hbβ subunit gene; Fig 1), while the Atlantic salmon has the highest number of hemoglobin subunit genes found among teleosts or gnathostomes (43 Hb genes in total; 20 of Hbα and 23 of Hbβ subunit) followed by the northern pike (*Esox lucius*) with 38 Hb genes (18+20). The Nile tilapia (*Oreochromis niloticus*), with the repertoire of 32 hemoglobin subunit genes (17 Hbα and 15 Hbβ subunits) has the highest number of hemoglobin genes among cichlids.

Most of the analyzed species possess two hemoglobin clusters, with the exception of the spotted gar (*Lepisosteus oculatus*) and the Australian ghostshark (*Callorhinchus milii*), which both have a single hemoglobin cluster (Figure 1). In addition, the northern pike (*Esox lucius*) and the Atlantic salmon (*Salmo salar*) with three and four hemoglobin clusters, respectively, demonstrate a variability in genomic organization of hemoglobin genes even within a single evolutionary lineage (Fig 1). The flanking genes upstream and downstream of each cluster further help to shed light on evolution of the whole paralogons. We found that synteny of the hemoglobin clusters and flanking genes is highly conserved across major evolutionary groups of teleosts, yet with certain deviations (Fig. 1).

### Evolutionary history of the hemoglobin subunit genes

In total, we identified and extracted the sequences of 197 hemoglobin alpha (Hbα) subunit genes and 185 hemoglobin beta (Hbβ) subunit genes from the 24 analyzed species (Table 1). We reconstructed the gene trees (Fig. 2; Fig. S1 and S2) and identified two major clades confirming the Hbα and Hbβ subunits in both approaches. For both Hbα and Hbβ subunit genes, the phylogenetic signal revealed several independent gene clades that have diversified in the teleost ancestor. Tetrapod genes correspondingly branch at the base of the tree, whereas teleost Hbα and Hbβ subunits are organized in four and three-to-four main clades each, respectively. The clades partially correspond to the genomic location either in LA or MN cluster, but not for all genes. The overall phylogenetic signal is more robust for the Hbβ subunit genes, while less clear for the Hbα subunit gene tree, which shows some polytomy and unresolved topology within one of the clades with mixed MN/LA genes (Fig. 2; Supp Fig S1 and S2). One of the clades within the Hbβ subunits is formed by predominantly cathodic subunits. Overall, the observed phylogenetic pattern suggests a presence of at least four putative copies of Hbα subunits and three or four Hbβ subunits in the ancestor of teleost fishes (Fig. 2).

**Figure 2:**
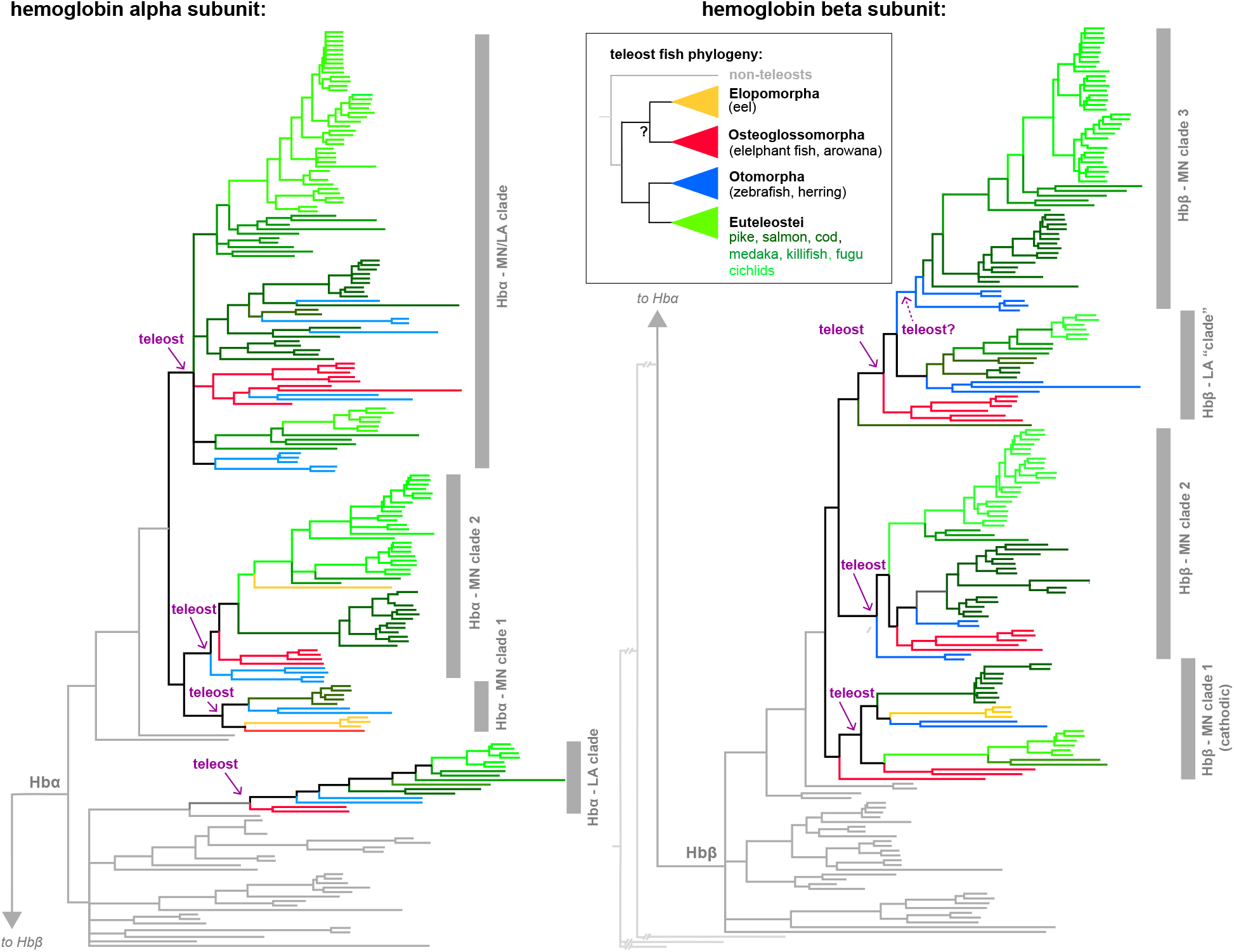
Gene trees of the Hbα and Hbβ hemoglobins subunits reconstructed by Bayesian analysis. Four main teleost lineages are highlighted by different colours (see inset for the teleost phylogeny) or black to elucidate Hb gene diversification in the teleost ancestor as well as after the teleost radiation. Non-teleost fish and tetrapod Hb genes are colored as grey. The tree topology suggests that the teleost ancestor had at least four Hbα genes and three (or four) Hbβ genes before teleost diversification corresponding to the recovered clades marked and labelled by purple arrow. Note that the duplication involving the LA clade Hbα genes and the rest Hbα genes possibly happened before teleost diversification (hence teleosts are found in two different clades of the tree). For the full-size gene trees with the copy names please refer to Supplementary Figures S1 and S2.

### Comparative genomics of hemoglobin clusters in cichlids

We have focused on the genomic architecture of the hemoglobin clusters in six cichlid species. The cichlid comparison revealed that Nile tilapia carries signals of numerous gene duplications of the two gene-rich regions within the MN cluster, one of the older and one of more recent evolutionary origins (Figure 3, Figure 4). The synteny in cichlids seems to be conserved among species, albeit not identical, suggesting ongoing genome rearrangements, as African and Neotropical cichlids differ slightly in the composition of their clusters (Figure 5, Supplementary Figure 5). Five of the high-quality genomes produced with the PacBio sequencing technology (*Amphilophus citrinellus*, *Archocentrus centrarchus*, *Astatotilapia calliptera*, *Maylandia zebra, Oreochromis niloticus;* retrieved from GenBank; Table 1) have been complemented by one species sequenced by the Oxford Nanopore (*Pungu maclareni*; this study). All cichlid species have two hemoglobin clusters (LA and MN). The LA cluster is conserved in all studied species and contains 3 hemoglobin genes in the same orientation flanked with lcmt1+aqp8-like+aqp8 upstream of the cluster and rhbdf1b+foxj1b+ubald1a in the downstream region (Fig 1). The MN cluster differs in number and orientation of hemoglobin subunits – from 15 hemoglobin subunit genes in *Amphilophus citrinellus*, 18 in *Maylandia zebra*, 19 in *Pungu maclareni*, 20 in *Archocentrus centrarchus*, 23 in *Astatotilapia calliptera* to, finally, *Oreochromis niloticus* with 29 hemoglobin genes, the species with the highest number so far identified among cichlids (Fig 1). Flanking genes in the MN cluster possess the same synteny – rhbdf1+mpg+nprl3 upstream from the cluster and kank2+dock6+elav13 from the downstream in all investigated cichlid species. Synteny plots on the MN cluster revealed certain rearrangements (Fig 3, 5, S5). While Nile tilapia and *Astatotilapia calliptera* carry two regions with high-similarity genes, *Maylandia zebra* and *Archocentrus centrarchus* contain only one of such high-similarity regions, and the Neotropical *Amphilophus citrinellus* has no high similarity regions (Supp Fig S4).

**Figure 3.**
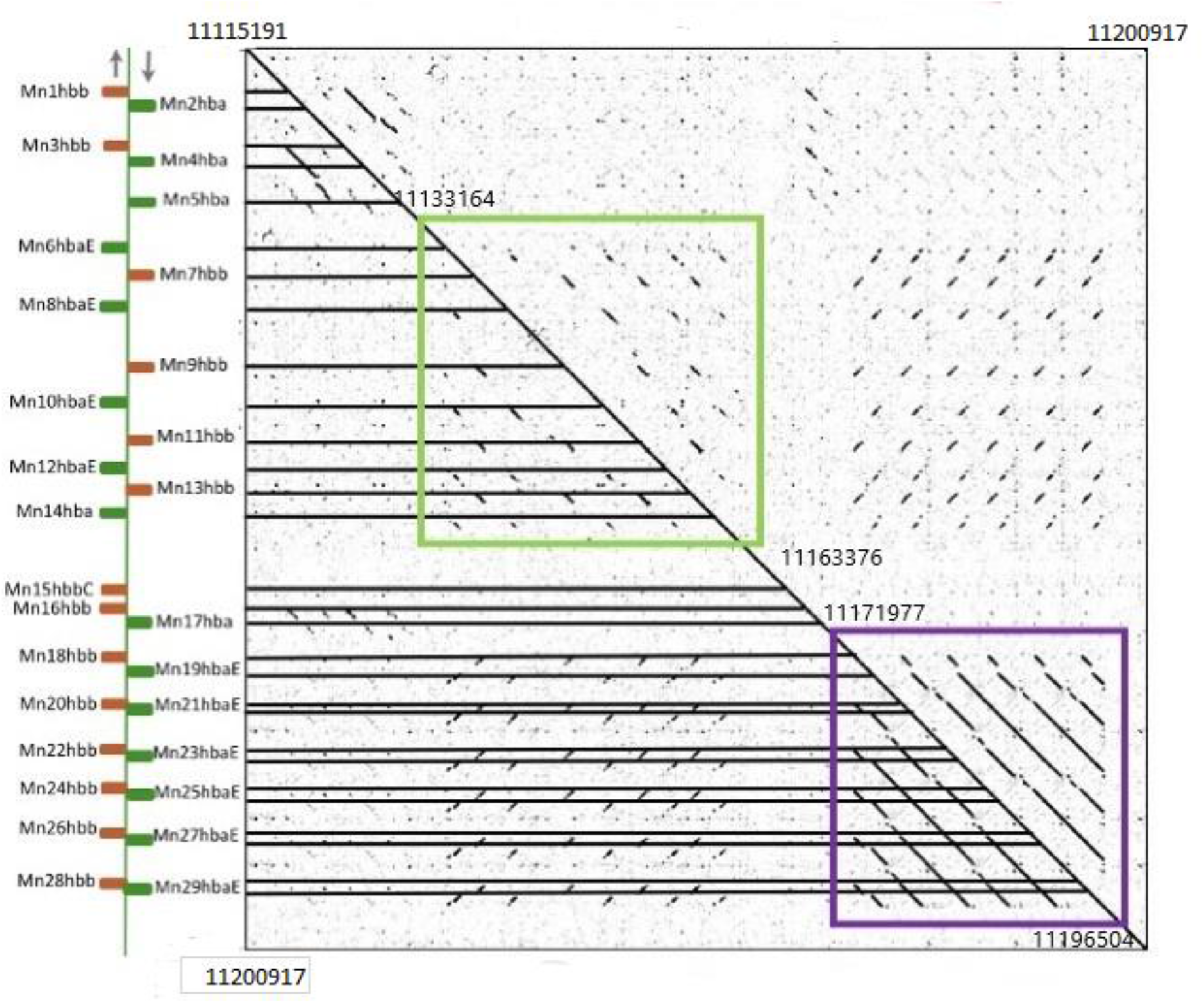
Gepard dotplot of the MN cluster of Nile tilapia (*Oreochromis niloticus*). The hemoglobin MN cluster is plotted against itself to reveal high-similarity regions (lines formed by black dots). Synteny of the hemoglobin genes in the cluster is shown on the left. Hbα subunit genes are colored green, while Hbβ subunits are marked in orange. Horizontal lines represent the first codon of each hemoglobin gene. There are two main regions within the MN cluster with reversed orientation to each other. The duplications within the region between positions 11’133’164 and 11’163’376 (marked by the green rectangle) are evolutionarily older, whereas the region between positions 11’171’977 and 11’196’504 (marked by the purple rectangle) is a result of more recent duplications (likely specific for tilapia or an oreochromine ancestor; Fig. 4). The suggested evolutionary scenario of hemoglobin gene evolution is presented in Figure 4. Genomic coordinates correspond to the reverse complemented genomic sequence (NCBI accession number NC_031969.2)

**Figure 4-.**
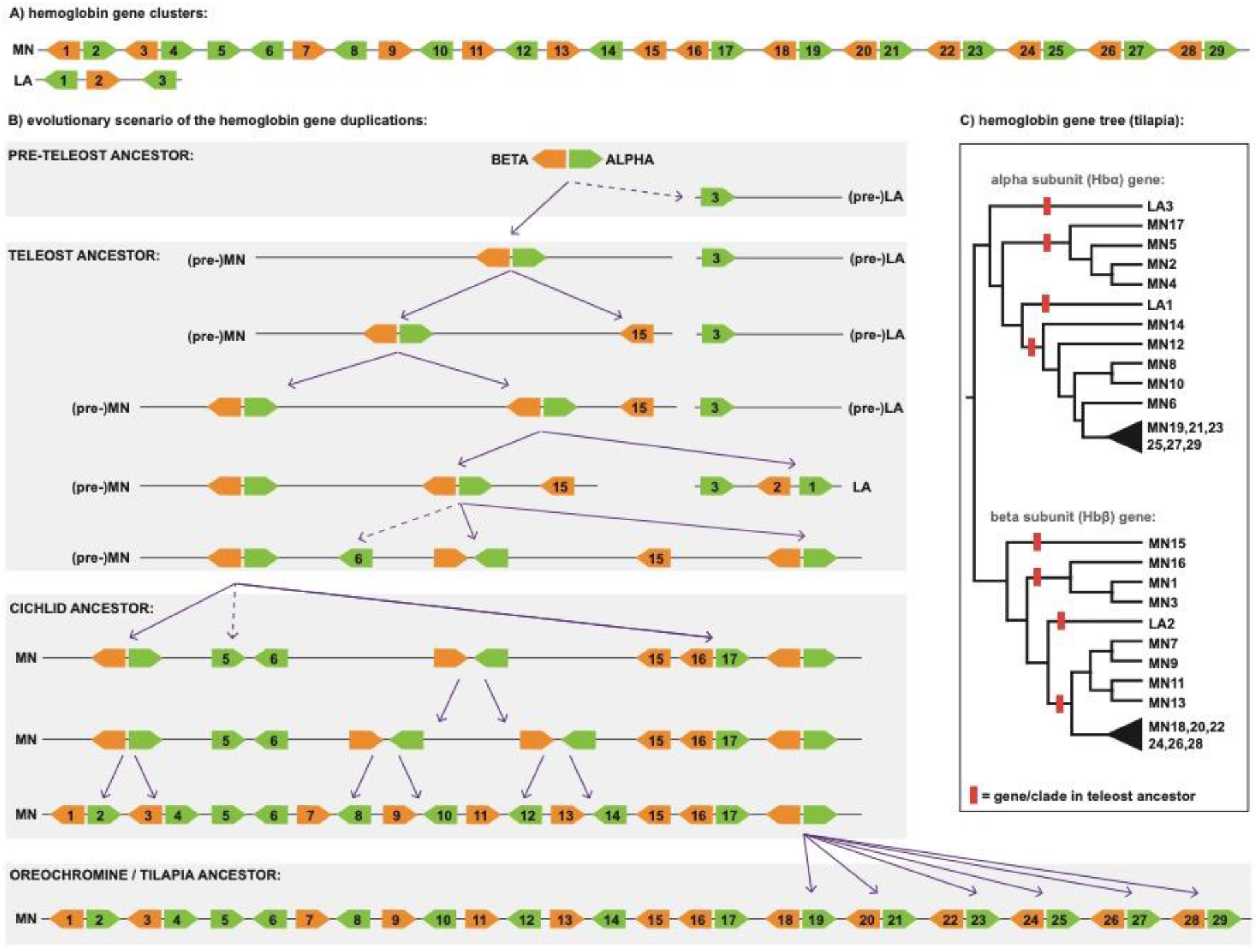
Evolutionary scenario of the Nile tilapia (*Oreochromis niloticus*) hemoglobin gene clusters. A) The conserved LA cluster is composed of three Hb genes, while the more dynamic MN cluster is composed of 29 Hb genes, the highest number among cichlids. B) Evolutionary scenario showing the series of events as reconstructed and interpreted from C) and Fig. 2, S1, S2, 3. The gene duplication marked by arrow. The gene numbers mark the first appearance of the recent Hb genes after which no more subsequent duplications have occurred. The main events occurred in the pre-teleost, teleost, cichlid, or oreochromine/*Oreochromis* ancestor (timing interpreted from the topology of the gene trees). Note that the genes currently located at the LA cluster have an older origin (pre-teleost and teleost), while the genes within the MN cluster proliferated by further gene duplications. C) the Nile tilapia Hb gene tree extracted from Fig 2 and Fig S1, S2. C) The hemoglobin gene tree of the Nile tilapia extracted from the overall gene tree (i.e., Hbα and Hbβ subunit gene trees in Fig. 2 and Supplementary Figure S1 and S2). Hbα subunit genes in green, Hbβ subunit genes in orange.

**Figure 5.**
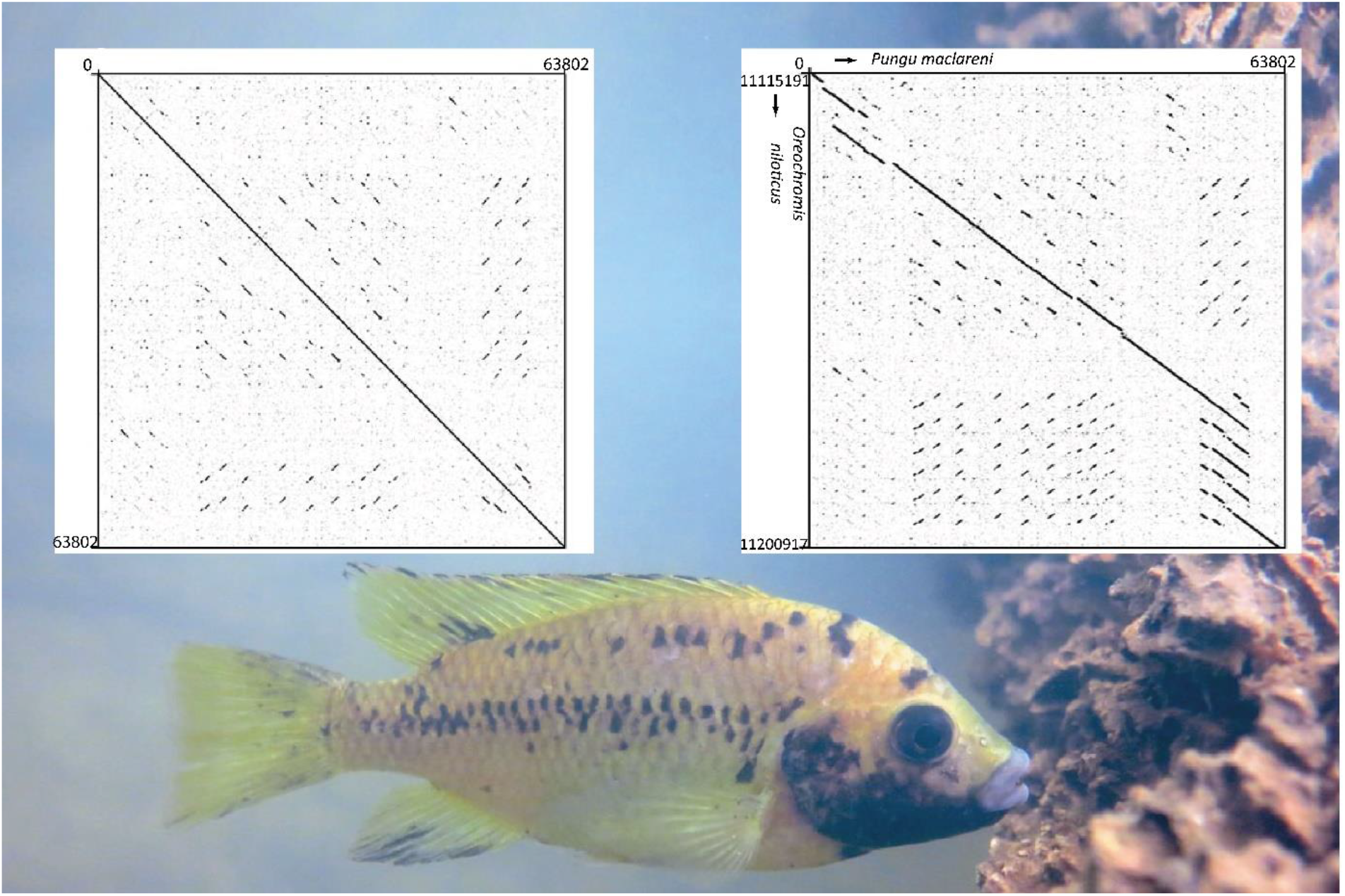
Comparison of the Nile tilapia and *Pungu maclareni* hemoglobin MN clusters. Gepard dotplot of the MN clusters from *Pungu maclareni* plotted against itself (left) and against the MN cluster of Nile tilapia (*Oreochromis niloticus*; right). Bottom: *Pungu maclareni*, the species from the Barombi Mbo crater lake sequenced within this study shown on the photo.

### Evolution of the hemoglobin repertoire in Nile Tilapia

The Nile tilapia (*Oreochromis niloticus*), similarly to other cichlids, has two hemoglobin clusters, the conserved LA cluster and the MN cluster, which demonstrates greater evolutionary dynamics. During evolution the ancestral Hbα and Hbβ subunit genes have diversified in teleosts. The evolutionarily oldest (and putatively functionally most divergent) are the cathodic Hbβ (MN15 in tilapia) and LA cluster Hbα (LA3) subunit genes, which have diversified in the teleost ancestor (or even earlier in case of LA)(Figure 4). Further, the hemoglobin cluster underwent downstream and upstream expansion in gene numbers because of the multiple tandem duplication events that happened in the teleost ancestor. Cichlids seem to further have diversified their hemoglobin gene repertoire namely in the MN cluster with many genes. Nile tilapia also carries traces of a recent gene expansion as noticed by the MN18 – 29 genes. Being partially shared with *Pungu maclareni* (Figure S1 and S2), these multiplications most likely started in the oreochromine ancestor (i.e. of both *Oreochromis* and *Pungu*) and further continued exclusively in *Oreochromis*.

### Positive selection test

We performed a test for positive selection on 186 and 175 full-length non-pseudogenized sequences of the Hbα and Hbβ hemoglobin subunit genes. Both Hbα and Hbβ subunits share similar pattern of selective pressures acting on them (Fig S5 and S6). We found an excess of non- synonymous mutations in the cichlid hemoglobin subunit genes that possibly appeared as a result of lineage-specific tandem duplications. This pattern can be observed for example in recently duplicated genes (Mn 1 Hbβ and Mn 3 Hbβ together with tandemly duplicated Mn 18 Hbβ, Mn 20 Hbβ, Mn 22 Hbβ, Mn 24 Hbβ, Mn 26 Hbβ and Mn 28 Hbβ *Oreochromis niloticus,* Mn 1 Hbβ, MN 2 Hbα and Mn 5 Hbβ, MN 6 Hbα in *Maylandia zebra*, MN 2 Hbα, MN4 Hbα, MN 6 Hbα and Mn 17 Hbβ, Mn 19 Hbβ and Mn 21 Hbβ in *Astatotilapia calliptera*, MN 7 Hbα and MN 11 Hbα, Mn 5 Hbα, Mn 6, Mn 8 Hbβ and Mn 18 Hbα, Mn 19 Hbα in *Archocentrus centrarchus,* Mn 16 Hbβ and Mn 18 Hbβ in *Pungu maclareni*), in which bursts of positive selection were detected prior to their emergence (Fig S5 and S6). In some cases, a similar patterns were also found in non-cichlids with expanded Hb gene repertoire, such as pike (De 1 Hbα and De 8 Hbα) and salmon (Mn 1.10 Hbα and Mn 2.13 Hbα). Signs of elevated positive selection is also found in the cathodic Hbβ subunit genes of African cichlids. (Fig S5 and S6)

## Discussion

Over evolutionary time, teleost fishes have undergone remarkable genetic changes in response to selective pressures associated with their aquatic habitats. Understanding the molecular evolution of fish hemoglobins not only elucidates the genetic basis for their functional diversity but also offers insights into interplay between genes, environment, and adaptation in the aquatic realm. In this study we applied comparative genomics to trace the evolutionary history of hemoglobin genes across different teleost (and namely cichlid) fish species, revealing namely patterns of gene duplications, losses, and genome rearrangements.

While hemoglobins are well studied and understood from many perspectives, the molecular basis and effects of single mutations are still less known than for example for rhodopsins (Yokoyama, 2008, Musilova et al., 2021). Major steps in evolution of hemoglobins, such as emergence of tetramers from the monomers and via dimers, has been described at the molecular level only recently (Pillai et al., 2020) by identification of the key amino acid substitutions, and this enables opportunities for subsequent research.

### Hemoglobin gene repertoire organized in clusters and the whole-genome duplication(s)

Teleost fishes exhibit an overwhelming morphological, ecological and taxonomical diversity, and, similarly, we also observe diversity in the genomic organization of the hemoglobin gene clusters. While the ancestral vertebrate whole-genome duplications played a crucial role in the origin of hemoglobins (Hoffman et al., 2012), we report that the hemoglobin diversity has been mostly driven by gene (mostly tandem) duplications contributing possibly to the adaptive potential of hemoglobins (Storz, 2016). The ancestor of jawed vertebrates most likely had a single hemoglobin gene pair, i.e., one Hbα and one Hbβ subunit gene (Fig. 4; Opazo et al., 2013). An organization of Hb genes in a single cluster is also found in a non-teleost ray-finned fish (i.e., gar – *Lepisosteus oculatus*) and in cartilaginous fishes (Fig. 1), while tetrapods and teleosts possess two clusters in their genome (most likely of independent origin since there are no similarities in the flanking genes nor the Hb gene sequence) (Fig. 1). The origin of the second, shorter, cluster in teleosts (labelled as LA cluster) has been previously associated with the teleost-specific whole genome duplication (TSGD) (Opazo et al., 2013). Our data does not clearly confirm this hypothesis. One of the Hbα genes (sometimes also referred to as Hbα subunits D-like due to their similarity to the hemoglobin D; Ahmed et al., 2020; Fig. 2, Suppl Fig S1) located in the LA cluster seems to have an older evolutionary origin, i.e., diversifying prior to the emergence of teleosts, [i.e., the teleost LA Hbα genes group together with one gar (non-teleost) Hbα gene, while other teleost Hbαs are more related to other two gar Hbα hemoglobins (Fig. 2, 4, S1)]. On the other hand, the paralogous flanking gene *rhbdf1* is found in both LA and MN clusters in all teleost lineages, suggesting the possible origin of the cluster by the TSGD. However, to test this we have reconstructed the gene tree of the *rhbdf1* genes from our data set (Figure S3) and found that the common origin of the two rhbdf1 paralogs is evolutionarily younger, in the Otomorpha + Euteleostei ancestor. There are clear gene clades of the LA and MN *rhbdf1* of otomorphs (e.g. zebrafish) + euteleosts (e.g. tilapia), while the elopomorph (eel) and osteoglossomorph (elephant fish) MN and LA copies form a separate clade (Figure S3). As a conclusion, we suggest that the evolution of the Hb genes most likely preceded their organization into clusters, i.e., some of the existent genes (of pre-teleost origin) have been reorganized into newly emerged clusters later (post-teleost diversification). This is also supported by the grouping of teleost LA Hbα genes with one gar Hbα gene in the gene tree (Fig S1), while the physical location of this gene in the gar genome is within a single gene cluster together with all other hemoglobins (Fig. 1). Interestingly, reorganization of Hb genes seems to be a common feature also among tetrapods, some of which have experienced massive hemoglobin gene reorganization into separate Hbα and Hbβ clusters (Fig. 1; Hardison, 2012). In sharks, gars and teleosts, on the other hand, the Hbα and Hbβ subunit genes are mixed in the hemoglobin clusters in the Hbα - Hbβ couples.

Teleosts seem to have the highest number of hemoglobin genes among vertebrates so far, ranging from five (fugu) to 43 found in salmon (Fig 1) compared to two hemoglobins in shark, seven in zebra finch, ten in mouse and human, and 16 in clawed frog. All teleosts have at least two hemoglobin clusters, LA and MN cluster. We found that northern pike and Atlantic salmon have an increased number of hemoglobin clusters and genes. Pike has 38 Hb genes located in three hemoglobin clusters (LA, MN and DE; the latter two on the same chromosome but in two separate regions). Salmon has 43 Hb genes in four hemoglobin clusters thanks to the *Salmoniformes*-specific whole genome duplication twelve million years ago (Koop et al., 2008). Interestingly, it has been shown that in salmon, some LA genes have an MN origin, bringing further evidence for common rearrangements of the Hb genes between clusters (Quinn et. al, 2010). Contrary to pike and salmon, *Tetraodontiformes* are known for an outstanding genome compaction (Brenner et al., 1993), which also affected hemoglobin genes in the MN cluster. Fugu has the lowest number of Hb genes (5) of all known teleosts, possessing only two hemoglobin subunit genes in the MN cluster (while the LA cluster remains conserved with three genes as in most teleosts).

Our findings for Hb gene numbers are most likely not the final ranges for teleosts given that we focused only on 24 selected vertebrate species. We believe that future research including more species may reveal even higher numbers of Hb copies. It has been impossible to infer correct genomic structure of complex loci, such as hemoglobin clusters without high-quality genome sequencing with reads long enough to cover the high similarity regions (e.g., Malinsky et al., 2018). The upcoming routine application of the high-quality genome sequencing will certainly lead to future findings regarding the hemoglobin gene clusters. This may be especially relevant for the species with high number of Hb genes. Noteworthy, while we analyzed the most recent version of the pike and salmon genomes (with 38 and 43 Hb genes respectively), there were only 29 and 33 identified genes in the previous versions of these genomes. On the other hand, we’re aware of the possible assembly errors and overestimation of the gene copies in the repetitive gene clusters (Ko et al., 2022). Lastly, we also notice that there might be intraspecific variability among different individuals of the same species, especially in the number of identical genes (such as Hb18-29) in tilapia. Albeit that, we believe our study still provides a valuable reference for future research, and general comparison of Hb gene repertoires, even though the exact numbers might vary or change in the future.

### Diversification of the hemoglobin genes in teleosts

The origin of Hbα and Hbβ hemoglobin subunit genes is a result of an ancient duplication event that likely happened ca 450 million years ago in the ancestor of gnathostomes vertebrates (Czelusniak et al., 1982). Our phylogeny confirms this, as Hbα and Hbβ subunits form two well supported clades in the gene tree (Fig. 2; Supplementary Fig S1, S2).

We found three major clades within the Hbβ subunit gene tree (Fig. 2, Figure S2) as in a previous study (Opazo et al., 2013). Unlike the Hbα subunits, all Hbβ subunit genes seem to diversify after the teleost ancestor (Fig 2, S2). The earliest branching genes are the cathodic hemoglobins (labelled as Hbβ MN15 in tilapia) located within the MN cluster. Cathodic hemoglobin gene (β subunit) is found as a single copy gene in most of the studied species, with the exception of pike (*Esox lucius*), Atlantic cod (*Gadus morhua*) and Japanese pufferfish (*Takifugu rubripes*) where it has been lost, and in European eel and Atlantic salmon, where it has been duplicated into multiple copies (Fig. S2).

The teleost ancestor had clearly at least four Hbα genes, whereas the signal is not that clear for Hbβ. The inferred phylogenetic pattern indicates the presence of at least three putative Hbβ copies in the teleost ancestor, which have emerged by gene duplications. Most likely, there was another duplication leading to four ancestral teleost Hbβ genes (and lost in some lineages), or that the last duplications happened later in evolution, in the ancestor of Otomorpha+Euteleostei. Our gene-trees (Fig. 2, Figure S2) suggest the latter, while the reconstructed scenario and the physical location in LA vs. MN clusters speaks rather for the former option. Additional analyses with more species are needed to resolve this uncertainty.

Our Hbα subunit gene tree is partially discordant with previously reported phylogenies, and the LA clade 2 (as in Opazo et al., 2013) could not be easily delineated; instead the genes are grouped within the combined MN/LA clade. Unlike most other clades within the gene tree, this clade is poorly supported, which could have been influenced by factors such as gene conversion or strong selection driving convergent evolution. Gene conversion and subsequent non-reciprocal exchange of the genetic material could in fact obscure the phylogenetic inference in paralogous genes (Archibald & Roger, 2002, Ratnakumar et al., 2010, Cortesi et al., 2015) although no conversion has been recently detected by a targeted analysis in ray-finned fish hemoglobin genes (Mao et al., 2023). Strong positive selection has also been found in the mixed Hbα clade (see below) and could partially explain the poor resolution of - and within - this clade (Figures S5, S6). These factors are, however, unlikely to compromise the overall strong phylogenetic signal, as we were generally able to recover well supported clades (with the aforementioned exception).

### Evolution of hemoglobins in cichlid fishes

Cichlids are characterized by a relatively high number of hemoglobin subunit genes compared to other studied teleosts (Fig. 1), ranging from 18 hemoglobin genes in *Amphilophus citrinellus* up to 32 in *Oreochromis niloticus.* Only two teleost species (pike, salmon) have a greater hemoglobin gene repertoire (38 and 43, respectively). All studied cichlids share the same synteny of the LA cluster including hemoglobins and three flanking genes upstream and downstream of the cluster. We report variation in the gene copies solely within the MN cluster of cichlids and this is probably a result of tandem duplications. Some Hb genes from African and Neotropical cichlids form sister clades suggesting emergence of many copies prior to cichlid diversification, whereas a substantial proportion of the gene diversity was apparently driven by lineage-specific or species-specific duplication events that happened after the split of the African and Neotropical lineages throughout cichlid evolution.

It has previously been shown that cichlids exhibit elevated number of duplicated genes in general (Brawand et al., 2014, Berner & Salzburger, 2015), and possibly this might have contributed to their evolutionary success – resulting in large numbers of species and adaptive radiations that enabled them to conquer various ecological and trophic niches. More specifically, Victoria cichlids are known to switch between hemoglobin types to cope with hypoxia (Rutjes et al., 2007; Thillart et al., 2018), and signs of positive selection have been reported in four hemoglobin genes from the MN cluster in deep-water Malawi cichlid from *Diplotaxodon* genus (Malinsky et al., 2017). Since hemoglobins are putative candidates for adaptive evolution, we perform the analysis of positive selection on the hemoglobin genes. We found an excess in non-synonymous-to synonymous mutations in some (but not all) duplicated genes of cichlid species, however the pattern is not clear. Most of the genes under positive selection result from recent duplications, might have experienced bursts of positive selection, namely the tilapia recent duplicates MN1 Hbβ and MN3 Hbβ subunit genes (with elevated positive selection signal). Interestingly, Nile tilapia itself is not a part of any cichlid radiation although it has the highest number of Hb genes so far. However, it is a species quite tolerant to hypoxia (Bergstedt et al., 2021), hence, there may be a biological relevance or traces of positive selection speaking for episodes of recent selection.

## Conclusion

We apply comparative genomics to reveal great diversity of the hemoglobin gene repertoires. Gene duplications, gene losses and genomic rearrangements have all contributed to the genomic architecture of hemoglobin subunit genes in teleost fishes. We conclude that tandem duplications played a major role in the dynamic increase of hemoglobin gene copies within the MN cluster of cichlid fishes. Furthermore, our results demonstrate that some genes withing the LA cluster had a pre-teleost origin, while the LA cluster itself may be evolutionarily younger, speaking for genome rearrangements. Contrarily, the MN cluster underwent very dynamic evolution – from a dramatic genome compaction in fugu with only two genes preserved in the MN cluster to the expansion in cichlid lineage (up to 29 genes), or furthermore, to duplication of both clusters in salmonid fishes. The teleost ancestor had at least four Hbα genes and three or four Hbβ genes, and the entire diversity of the hemoglobin gene repertoires has arisen from these ancestral copies. This study aims to serve as a valuable overview for the future research aiming to integrate observation from the wild with physiological experiments, targeted mutagenesis and in vitro protein engineering to understand the mechanisms of hemoglobin molecular evolution.

## Acknowledgements

We would like to express our thanks to Veronika Truhlarova for a technical support, Helle Tessand Baalsrud, Sissel Jentoft and Monica Hongrø Solbakken for the help with the entire first steps in the bioinformatic analyses, Demian Burguera for his bioinformatic insight, Daniel Elleder for help with the long-fragment DNA extraction and Petr Pajer for his valuable help and performing the Nanopore sequencing protocol. We further thank local people of the Barombi village to allow us to fish in the Barombi Mbo lake, the Ministry of Scientific Research and Innovation in Cameroon to grant us research permits (for D.O., A.I., A.R.B.N., Z.M.). Special thanks to Herman Igor Kittio for help with the fieldwork. Computational resources were supplied by the project “e-Infrastruktura CZ” (e-INFRA CZ LM2018140) supported by the Ministry of Education, Youth and Sports of the Czech Republic. The project was funded by the Czech Science Foundation (21-31712S) and Charles University Grant Agency (GAUK 1556119).

## Supplementary

separate file

## Data availability

Sequence alignment of all hemoglobin genes in FASTA format. Set of genomic clusters with the final annotation (as Geneious, or exported as .gb file).

## References

Ahmed, M. H., Ghatge, M. S., & Safo, M. K. (2020). Hemoglobin: structure, function and allostery. Vertebrate and invertebrate respiratory proteins, lipoproteins and other body fluid proteins, 345–382.

Archibald, J. M., & Roger, A. J. (2002). Gene conversion and the evolution of euryarchaeal chaperonins: a maximum likelihood-based method for detecting conflicting phylogenetic signals. Journal of Molecular Evolution, 55(2), 232–245.

Bergstedt, J. H., Pfalzgraff, T., & Skov, P. V. (2021). Hypoxia tolerance and metabolic coping strategies in Oreochromis niloticus. Comparative Biochemistry and Physiology Part A: Molecular & Integrative Physiology, 257, 110956.

Berner, D., & Salzburger, W. (2015). The genomics of organismal diversification illuminated by adaptive radiations. Trends in Genetics, 31(9), 491–499.

Bolognini, D., Bartalucci, N., Mingrino, A., Vannucchi, A. M., & Magi, A. (2019). NanoR: A user-friendly R package to analyze and compare nanopore sequencing data. PloS one, 14(5), e0216471.

Borza, T., Stone, C., Gamperl, A. K., & Bowman, S. (2009). Atlantic cod (*Gadus morhua*) hemoglobin genes: multiplicity and polymorphism. BMC genetics, 10(1), 1–14.

Brenner, S., Elgar, G., Sanford, R., Macrae, A., Venkatesh, B., & Aparicio, S. (1993). Characterization of the pufferfish (Fugu) genome as a compact model vertebrate genome. Nature, 366(6452), 265.

Broughton, R. E., Betancur-R, R., Li, C., Arratia, G., & Ortí, G. (2013). Multi-locus phylogenetic analysis reveals the pattern and tempo of bony fish evolution. PLoS currents, 5.

Chan, F. Y., Robinson, J., Brownlie, A., Shivdasani, R. A., Donovan, A., Brugnara, C., … & Zon, L. I. (1997). Characterization of adult α-and β-globin genes in the zebrafish. Blood, 89(2), 688–700.

Chen, Z., Qiao, F., He, Y., Xie, H., & Qi, D. (2016). Evidence for positive selection on α and β globin genes in pikas and zokor from the Qinghai-Tibetan Plateau. Gene & Translational Bioinformatics, 2.

Cortesi, F., Musilová, Z., Stieb, S. M., Hart, N. S., Siebeck, U. E., Malmstrøm, M., … & Salzburger, W. (2015). Ancestral duplications and highly dynamic opsin gene evolution in percomorph fishes. Proceedings of the National Academy of Sciences, 112(5), 1493–1498.

Czelusniak, J., Goodman, M., Hewett-Emmett, D., Weiss, M. L., Venta, P. J., & Tashian, R. E. (1982). Phylogenetic origins and adaptive evolution of avian and mammalian haemoglobin genes. Nature, 298(5871), 297.

Darriba, D., Taboada, G. L., Doallo, R., & Posada, D. (2012). jModelTest 2: more models, new heuristics and parallel computing. Nature methods, 9(8), 772.

Dehal, P., & Boore, J. L. (2005). Two rounds of whole genome duplication in the ancestral vertebrate. PLoS biology, 3(10), e314.

Di Prisco, G., Giardina, B., & Weber, R. E. (Eds.). (2000). Hemoglobin function in vertebrates: molecular adaptation in extreme and temperate environments. Springer Science & Business Media.

Fawcett, J. A., Maere, S., & Van De Peer, Y. (2009). Plants with double genomes might have had a better chance to survive the Cretaceous–Tertiary extinction event. Proceedings of the National Academy of Sciences, 106(14), 5737–5742.

Feng, J., Liu, S., Wang, X., Wang, R., Zhang, J., Jiang, Y., … & Liu, Z. (2014). Channel catfish hemoglobin genes: identification, phylogenetic and syntenic analysis, and specific induction in response to heat stress. Comparative Biochemistry and Physiology Part D: Genomics and Proteomics, 9, 11–22.

Garcia-Fernàndez, J. (2005). The genesis and evolution of homeobox gene clusters. Nature Reviews Genetics, 6(12), 881–892.

Gillemans, N., McMorrow, T., Tewari, R., Wai, A. W., Burgtorf, C., Drabek, D., … & Grosveld, F. (2003). Functional and comparative analysis of globin loci in pufferfish and humans. Blood, 101(7), 2842–2849.

Glasauer, S. M., & Neuhauss, S. C. (2014). Whole-genome duplication in teleost fishes and its evolutionary consequences. Molecular genetics and genomics, 289(6), 1045–1060.

Gillespie, John H. The causes of molecular evolution. Vol. 2. Oxford University Press on Demand, 1994.

Green, J., Corbet, S. A. & Betney, E. (1973) Ecological studies on crater lakes in West Cameroon The blood of endemic cichlids in Barombi Mbo in relation to stratification and their feeding habits. J Zool 170, 299–308.

Hahn, C., Genner, M. J., Turner, G. F., & Joyce, D. A. (2017). The genomic basis of cichlid fish adaptation within the deepwater “twilight zone” of Lake Malawi. Evolution letters, 1(4), 184–198.

Hardison, R. C. (2008). Globin genes on the move. Journal of biology, 7(9), 35.

Hardison, R. C. (2012). Evolution of hemoglobin and its genes. Cold Spring Harbor perspectives in medicine, 2(12), a011627.

Hoang, D. T., Chernomor, O., Von Haeseler, A., Minh, B. Q., & Vinh, L. S. (2017). UFBoot2: improving the ultrafast bootstrap approximation. Molecular biology and evolution, 35(2), 518–522.

Holland, P. W., Garcia-Fernàndez, J., Williams, N. A., & Sidow, A. (1994). Gene duplications and the origins of vertebrate development. Development, 1994(Supplement), 125–133

Hoffmann, F. G., Opazo, J. C., & Storz, J. F. (2012). Whole-genome duplications spurred the functional diversification of the globin gene superfamily in vertebrates. Molecular biology and evolution, 29(1), 303–312.

Hughes, A. L., & Friedman, R. (2003). 2R or not 2R: testing hypotheses of genome duplication in early vertebrates. In Genome Evolution (pp. 85–93). Springer, Dordrecht.

Hurley, I. A., Mueller, R. L., Dunn, K. A., Schmidt, E. J., Friedman, M., Ho, R. K., … & Coates, M. I. (2006). A new time-scale for ray-finned fish evolution. Proceedings of the Royal Society B: Biological Sciences, 274(1609), 489–498.

Jaillon, O., Aury, J. M., Brunet, F., Petit, J. L., Stange-Thomann, N., Mauceli, E., … & Nicaud, S. (2004). Genome duplication in the teleost fish Tetraodon nigroviridis reveals the early vertebrate proto-karyotype. Nature, 431(7011), 946.

Jansen, H. J., Liem, M., Jong-Raadsen, S. A., Dufour, S., Weltzien, F. A., Swinkels, W., … & Van den Thillart, G. E. (2017). Rapid de novo assembly of the European eel genome from nanopore sequencing reads. Scientific reports, 7(1), 1–13

Jeffreys, A. J., Wilson, V., Wood, D., Simons, J. P., Kay, R. M., & Williams, J. G. (1980). Linkage of adult α-and β-globin genes in X. laevis and gene duplication by tetraploidization. Cell, 21(2), 555–564.

Kalyaanamoorthy, S., Minh, B. Q., Wong, T. K., von Haeseler, A., & Jermiin, L. S. (2017). ModelFinder: fast model selection for accurate phylogenetic estimates. Nature methods, 14(6), 587.

Kasahara, M., Naruse, K., Sasaki, S., Nakatani, Y., Qu, W., Ahsan, B., … & Jindo, T. (2007). The medaka draft genome and insights into vertebrate genome evolution. Nature, 447(7145), 714.

Katoh, K., Misawa, K., Kuma, K. I., & Miyata, T. (2002). MAFFT: a novel method for rapid multiple sequence alignment based on fast Fourier transform. Nucleic acids research, 30(14), 3059–3066.

Katoh, K., & Standley, D. M. (2013). MAFFT multiple sequence alignment software version 7: improvements in performance and usability. Molecular biology and evolution, 30(4), 772–780.

Kautt, A. F., Machado-Schiaffino, G., & Meyer, A. (2016). Multispecies outcomes of sympatric speciation after admixture with the source population in two radiations of Nicaraguan crater lake cichlids. PLoS genetics, 12(6), e1006157.

Ko, B. J., Lee, C., Kim, J., Rhie, A., Yoo, D. A., Howe, K., … & Kim, H. (2022). Widespread false gene gains caused by duplication errors in genome assemblies. Genome Biology, 23(1), 205.

Koop, B. F., Von Schalburg, K. R., Leong, J., Walker, N., Lieph, R., Cooper, G. A., … & Brahmbhatt, S. (2008). A salmonid EST genomic study: genes, duplications, phylogeny and microarrays. BMC genomics, 9(1), 545.

Krumsiek, J., Arnold, R., & Rattei, T. (2007). Gepard: a rapid and sensitive tool for creating dotplots on genome scale. Bioinformatics, 23(8), 1026–1028.

Lee, J. Y., Kong, M., Oh, J., Lim, J., Chung, S. H., Kim, J. M., … & Kwak, W. (2021). Comparative evaluation of Nanopore polishing tools for microbial genome assembly and polishing strategies for downstream analysis. Scientific Reports, 11(1), 20740.

Lei, Y., Yang, L., Zhou, Y., Wang, C., Lv, W., Li, L., & He, S. (2021). Hb adaptation to hypoxia in high-altitude fishes: Fresh evidence from schizothoracinae fishes in the Qinghai-TiHbβn Plateau. International Journal of Biological Macromolecules, 185, 471–484.

Makałowski, W. (2001). Are we polyploids? A brief history of one hypothesis. Genome Research, 11(5), 667–670.

Malinsky, M., Challis, R., Tyers, A. M., Schiffels, S., Terai, Y., Ngatunga, B. P., Miska, E. A., Durbin, R., Genner, M. J., Turner, G. F. (2015). Genomic islands of speciation separate cichlid ecomorphs in an East African crater lake. Science (New York, N.Y.), 350(6267), 1493–8.

Malinsky, M., Svardal, H., Tyers, A.M., Genner, M.J., Turner, G.F., Miska, E.A. & Durbin, R. (2018) Whole-genome sequences of Malawi cichlids reveal multiple radiations interconnected by gene flow. Nature Ecology & Evolution 2, 1940–1955.

Marino, K., Boschetto, L., de Pascale, D., & Cocca, E. (2007). Organisation of the Hb 1 genes of the Antarctic skate Bathyraja eatonii: New insights into the evolution of globin genes. Gene, 406(1- 2), 199–208.

McKiernan, H. E., & Danielson, P. B. (2017). Molecular diagnostic applications in forensic science. In Molecular diagnostics (pp. 371-394). Academic Press.

Miller, M. A., Pfeiffer, W., & Schwartz, T. (2010, November). Creating the CIPRES Science Gateway for inference of large phylogenetic trees. In 2010 gateway computing environments workshop (GCE) (pp.1-8).Ieee.

Moriyama, Y., & Koshiba-Takeuchi, K. (2018). Significance of whole-genome duplications on the emergence of evolutionary novelties. Briefings in functional genomics, 17(5), 329–338.

Mukherjee, S., Bartoš, O., Zdeňková, K., Hanák, P., Horká, P., & Musilova, Z. (2021). Evolution of the parvalbumin genes in teleost fishes after the whole-genome duplication. Fishes, 6(4), 70.

Murrell, B., Weaver, S., Smith, M. D., Wertheim, J. O., Murrell, S., Aylward, A., … & Scheffler, K. (2015). Gene-wide identification of episodic selection. Molecular biology and evolution, 32(5), 1365–1371.

Musilova, Z., Salzburger, W., & Cortesi, F. (2021). The visual opsin gene repertoires of teleost fishes: evolution, ecology, and function. Annual review of cell and developmental biology, 37, 441–468.

Nery, M. F., Arroyo, J. I., & Opazo, J. C. (2013). Genomic organization and differential signature of positive selection in the Hbα and Hbβ globin gene clusters in two cetacean species. Genome biology and evolution, 5(12), 2359–2367.

Nguyen, L. T., Schmidt, H. A., von Haeseler, A., & Minh, B. Q. (2014). IQ-TREE: a fast and effective stochastic algorithm for estimating maximum-likelihood phylogenies. Molecular biology and evolution, 32(1), 268–274.

Ohno, S. (2013). Evolution by gene duplication. Springer Science & Business Media.

Opazo, J. C., Butts, G. T., Nery, M. F., Storz, J. F., & Hoffmann, F. G. (2013). Whole-genome duplication and the functional diversification of teleost fish hemoglobins. Molecular biology and evolution, 30(1), 140–153.

Parey, E., Louis, A., Montfort, J., Guiguen, Y., Crollius, H. R., & Berthelot, C. (2022). An atlas of fish genome evolution reveals delayed rediploidization following the teleost whole-genome duplication. Genome Research, 32(9), 1685–1697.

Patel, V. S., Cooper, S. J., Deakin, J. E., Fulton, B., Graves, T., Warren, W. C., … & Graves, J. A. (2008). Platypus globin genes and flanking loci suggest a new insertional model for Hbβ-globin evolution in birds and mammals. BMC biology, 6(1), 34.

Philipsen, S., & Hardison, R. C. (2018). Evolution of hemoglobin loci and their regulatory elements. *Blood Cells*, Molecules, and Diseases, 70, 2–12.

Pillai, A. S., Chandler, S. A., Liu, Y., Signore, A. V., Cortez-Romero, C. R., Benesch, J. L., … & Thornton, J. W. (2020). Origin of complexity in haemoglobin evolution. Nature, 581(7809), 480–485.

Putnam, N. H., Butts, T., Ferrier, D. E., Furlong, R. F., Hellsten, U., Kawashima, T., … & Benito-Gutierrez, E. (2008). The amphioxus genome and the evolution of the chordate karyotype. Nature, 453(7198), 1064.

Ratnakumar, A., Mousset, S., Glémin, S., Berglund, J., Galtier, N., Duret, L., & Webster, M. T. (2010). Detecting positive selection within genomes: the problem of biased gene conversion. Philosophical Transactions of the Royal Society B: Biological Sciences, 365(1552), 2571–2580.

Rennison, Diana J., Gregory L. Owens, and John S. Taylor. “Opsin gene duplication and divergence in ray-finned fish.” Molecular phylogenetics and evolution 62, no. 3 (2012): 986–1008.

Ronald, J., & Akey, J. M. (2005). Genome-wide scans for loci under selection in humans. Human genomics, 2(2), 1–13.

Ronco, F., Matschiner, M., Böhne, A., Boila, A., Büscher, H. H., El Taher, A., … & Salzburger, W. (2021). Drivers and dynamics of a massive adaptive radiation in cichlid fishes. Nature, 589(7840), 76–81

Ronquist, F., Teslenko, M., Van Der Mark, P., Ayres, D. L., Darling, A., Höhna, S., … & Huelsenbeck, J. P. (2012). MrBayes 3.2: efficient Bayesian phylogenetic inference and model choice across a large model space. Systematic biology, 61(3), 539–542.

Rummer, J. L., McKenzie, D. J., Innocenti, A., Supuran, C. T., & Brauner, C. J. (2013). Root effect hemoglobin may have evolved to enhance general tissue oxygen delivery. Science, 340(6138), 1327–1329.

Rutjes, H. A., Nieveen, M. C., Weber, R. E., Witte, F., & Van den Thillart, G. E. E. J. M. (2007). Multiple strategies of Lake Victoria cichlids to cope with lifelong hypoxia include hemoglobin switching. American Journal of Physiology-Regulatory, Integrative and Comparative Physiology, 293(3), R1376–R1383.

Salzburger, W. (2018). Understanding explosive diversification through cichlid fish genomics. Nature Reviews Genetics, 19(11), 705–717.

Skrabanek, L., & Wolfe, K. H. (1998). Eukaryote genome duplication-where’s the evidence?. Current opinion in genetics & development, 8(6), 694–700.

Simão, F. A., Waterhouse, R. M., Ioannidis, P., Kriventseva, E. V., & Zdobnov, E. M. (2015). BUSCO: assessing genome assembly and annotation completeness with single-copy orthologs. Bioinformatics, 31(19), 3210–3212.

Stanke, M., & Morgenstern, B. (2005). AUGUSTUS: a web server for gene prediction in eukaryotes that allows user-defined constraints. Nucleic acids research, 33(suppl_2), W465–W467.

Steinke, D., Salzburger, W., Braasch, I., & Meyer, A. (2006). Many genes in fish have species-specific asymmetric rates of molecular evolution. BMC genomics, 7, 1–18.

Storz, J. F., Opazo, J. C., & Hoffmann, F. G. (2013). Gene duplication, genome duplication, and the functional diversification of vertebrate globins. Molecular phylogenetics and evolution, 66(2), 469–478.

Storz, J. F. (2016). Gene duplication and evolutionary innovations in hemoglobin-oxygen transport. Physiology, 31(3), 223–232.

Storz, J. F. (2018). Hemoglobin: insights into protein structure, function, and evolution. Oxford University Press.

Suyama, M., Torrents, D., & Bork, P. (2006). PAL2NAL: robust conversion of protein sequence alignments into the corresponding codon alignments. Nucleic acids research, 34(suppl_2), W609–W612.

Svardal, H., Salzburger, W. & Malinsky, M. (2021) Genetic Variation and Hybridization in Evolutionary Radiations of Cichlid Fishes. Annu Rev Anim Biosci 9, 55–79.

Team, R. C. (2018). A language and environment for statistical computing. R Foundation for Statistical Computing, Vienna, Austria. Available online: www.R-project.org/(accessed on 11 September 2020).

Van de Peer, Y., Maere, S., & Meyer, A. (2010). 2R or not 2R is not the question anymore. Nature Reviews Genetics, 11(2), 166–166.

van den Thillart, G., Wilms, I., Nieveen, M., Weber, R. E., & Witte, F. (2018). Hypoxia-induced changes in hemoglobins of Lake Victoria cichlids. Journal of Experimental Biology, 221(17), jeb177832.

Weber, R. E., & Jensen, F. B. (1988). Functional adaptations in hemoglobins from ectothermic vertebrates. Annual Review of Physiology, 50(1), 161–179.

Weber, R. E. (1990). Functional significance and structural basis of multiple hemoglobins with special reference to ectothermic vertebrates.: In: Animal Nutrition and Transport Processes. 2. Transport, Respiration and Excretion: Comparative and Environmental Aspects. Comparative Physiology (basel: Karger), 6, 58-75.

Wells, R. M. (1999). Evolution of haemoglobin function: molecular adaptations to environment. Clinical and Experimental Pharmacology and Physiology, 26(8), 591–595.

Wells, R. M. (2009). Blood-gas transport and hemoglobin function: Adaptations for functional and environmental hypoxia. In Fish physiology (Vol. 27, pp. 255-299). Academic Press.

Yang, Z. (2007). PAML 4: phylogenetic analysis by maximum likelihood. Molecular biology and evolution, 24(8), 1586–1591.

Yokoyama, S. (2008). Evolution of dim-light and color vision pigments. Annu. Rev. Genomics Hum. Genet., 9, 259–282.

Walker, B. J., Abeel, T., Shea, T., Priest, M., Abouelliel, A., Sakthikumar, S., … & Earl, A. M. (2014). Pilon: an integrated tool for comprehensive microbial variant detection and genome assembly improvement. PloS one, 9(11), e112963.

Wang, L. G., Lam, T. T. Y., Xu, S., Dai, Z., Zhou, L., Feng, T., … & Yu, G. (2020). Treeio: an R package for phylogenetic tree input and output with richly annotated and associated data. Molecular biology and evolution, 37(2), 599–603.

Wick, R. R., Judd, L. M., & Holt, K. E. (2019). Performance of neural network basecalling tools for Oxford Nanopore sequencing. Genome biology, 20, 1–10.

